# PIEZO1 upregulation in spinal cord astrocytes during MOG_35-55_-induced EAE correlates with ECM remodeling

**DOI:** 10.64898/2026.06.23.734091

**Authors:** Maik Hintze, Rittika Chunder, Michael Schwarz, Zafar Nurmatov, Markus Lorke, Jolanda Bäcker, Kim Leonie Holzbauer, Emily Brockmann, Arif Ekici, Aldo Roberto Boccaccini, Stefanie Kuerten

## Abstract

**Background:** Extracellular matrix (ECM) remodeling is increasingly recognized as an important component of neuroinflammatory pathology in multiple sclerosis (MS), yet the mechanisms by which CNS cells sense and respond to alterations in their mechanical environment and the spatial across which mechanical changes can influence cellular behavior remain poorly understood. Piezo1 is a mechanosensitive ion channel that regulates cellular responses to mechanical stimuli and has recently emerged as a potential modulator of neuroinflammation.

**Methods:** Experimental autoimmune encephalomyelitis (EAE) was induced in C57BL/6 wildtype mice using myelin oligodendrocyte glycoprotein (MOG):_35-55_. Immunohistochemical analyses were performed in spinal cord gray matter (GM), normal-appearing white matter (NAWM), and white matter lesion (LES) regions to assess ECM remodeling, total Piezo1 expression, and astrocyte-specific Piezo1 expression during acute and chronic EAE stages. Correlations with clinical EAE severity were determined. In parallel, mixed primary murine glial cultures were exposed to substrates of different stiffness and analyzed by transcriptomic profiling to investigate mechanobiological responses *in vitro*.

**Results:** ECM-associated proteins, including glial fibrillary acidic protein (GFAP), fibronectin-1 and matrix metalloproteinase-3 (MMP3), were regionally upregulated during EAE, indicating widespread tissue remodeling beyond focal inflammatory lesions. Total Piezo1 expression was increased within lesions and transiently elevated in GM, whereas astrocyte-specific Piezo1 remained persistently upregulated during both acute and chronic EAE. Astrocytic Piezo1 expression correlated closely with ECM remodeling and clinical EAE severity, particularly in GM and NAWM. Notably, both total and astrocyte-specific Piezo1 showed stronger associations with clinical disability than classical inflammatory markers. Transcriptomic analysis revealed pronounced stiffness-dependent responses in glial cells, including alterations in extracellular matrix organization, cytokine signaling, cell adhesion, and proliferative pathways.

**Conclusions:** Our findings identify astrocytic Piezo1 as a prominent component of neuroinflammatory tissue remodeling during EAE. The close association of Piezo1 with ECM alterations, clinical disease severity, and stiffness-dependent glial responses supports a link between neuroinflammation and mechanosensory signaling. These results highlight mechanosensation as a potentially important contributor to CNS pathology and establish Piezo1 alteration as a candidate biomarker for neuroinflammatory disease.

## Background

Multiple sclerosis (MS) is a chronic neuroinflammatory disease characterized by the recruitment of immune cells into the central nervous system (CNS), activation of microglia and astrocytes, axonal loss and oligodendrocyte death (Ghorbani et al., 2021). The CNS is composed of tightly packed neuronal and glial cells with comparatively little extracellular volume and hence, the mechanical properties of individual cells contribute substantially to general tissue mechanics (Pillai and Franze, 2024). The soft mechanical signature of astrocytes in comparison to neurons (Lu et al., 2006) therefore causes relative softness of glial scars after neuronal degeneration in the CNS (Moeendarbary et al., 2017).

Alteration of the extracellular matrix (ECM) is integral to MS pathology and strongly influences both injury and repair processes at all stages of the disease (Sobel et al., 2001), which is associated with diffuse brain tissue softening *in vivo* (Silva et al., 2024; Wuerfel et al., 2010). The ECM is composed of a mix of proteins and carbohydrates (Soles et al., 2023) which together form a dynamic and complex scaffold that not only provide biomechanical support for neurons and glial cells (van Horssen et al., 2007) but also play a key role in regulating ion homeostasis, cell signaling and synaptic plasticity (Ge et al., 2026). The functional attributes of the ECM arise from their composition and localization in different CNS compartments (Lau et al., 2013). For example, neural interstitial matrix components, like hyaluronan and proteoglycans, fill and structure the space between neurons and glial cells and therefore act as a biophysical lattice for the brain microenvironment (Tewari et al., 2022). The basement membrane ECM, primarily composed of fibrous proteins including collagens and laminins (Hallmann et al., 2020), provides structural support to the neurovascular unit and forms a biochemical barrier regulating vascular integrity (Reed et al., 2019).

Astrocytes have long been known to be major producers of both brain interstitial as well as basement membrane ECM (Wiese et al., 2012). Furthermore, ECM components secreted by astrocytes into the extracellular matrix can exert autocrine effects on their cellular behaviour, phenotype and gene expression profile (Baeten and Akassoglou, 2011; Hu et al., 2021; George and Geller, 2018). In the event of an inflammatory insult, the delicate bidirectional interaction between the ECM and astrocytes is severely disrupted (Woo and Sontheimer, 2023), causing these cells, which both secrete and respond to the ECM, to become reactive and thereby drive pathological ECM remodeling and altered mechanosensing (Moeendarbary et al., 2017; Berdiaki et al., 2024; Mierke et al., 2024).

Expression of both CNS parenchymal and basement membrane-related ECM proteins have been shown to be altered over the disease course in experimental autoimmune encephalomyelitis (EAE), a commonly used rodent model of MS (Mohan et al., 2010). At disease onset and peak, elevation of ECM proteoglycan in the white matter of the spinal cord has been shown to be closely associated with activated microglia, astrocytes and invading immune cells (Ghorbani et al., 2022). Similarly, glial scar formation, a chronic pathological process in EAE, is characterized by reactive astrocytes that drastically increase their synthesis of ECM components, such as chondroitin sulfate proteoglycans (CSPGs) and tenascins, which then accumulate in gliotic areas (Yahn et al., 2020; Tatomir et al., 2018).

These alterations in the ECM component abundance, or changes in ECM “stiffness”, profoundly influence cellular mechanotransduction (Berdiaki et al., 2024; Di et al., 2023). At the molecular level, mechanotransduction can trigger downstream signaling cascades associated with protein translation and cytokine secretion (Rocha et al., 2022), often through the activation of mechanosensitive ion channels. Although several CNS-expressing ion channels, including transient receptor potential vanilloid 4 (TRPV4) channels (Harraz et al., 2022) and potassium channel subfamily K member 2 (TREK1) (Hervieu et al., 2001), have been implicated in mechanosensation, only Piezo proteins, which are large trimeric transmembrane complexes (Wang et al., 2019; Saotome et al., 2018), are *bona fide* mechanically activated ion channels (Murthy et al., 2017; Coste et al., 2010).

In vertebrates, the evolutionarily conserved Piezo family of proteins comprises two homologs, Piezo1 and Piezo2 (Yuan et al., 2026). Piezo2 expression is primarily restricted to peripheral tissues, including the sensory ganglia (Zhang et al., 2026), respiratory organs, and gastrointestinal tract (Servin-Vences et al., 2023). Piezo1 is broadly expressed across a variety of immune cells and in the CNS by both neuronal and non-neuronal cells (Cudmore and Santana, 2022; Chi et al., 2022; Zhu et al., 2023), implying an important role in mechano-immune regulation in CNS diseases (Yang et al., 2025; Ikiz et al., 2024).

One study, for example, has shown that deletion of Piezo1 in T cells reduces EAE severity (Jairaman et al., 2021), thereby highlighting that dysregulation – whether through an increase or decrease in Piezo1 activity or expression – can disrupt cellular homeostasis and contribute to the development and progression of disease pathogenesis (Zong et al., 2023; Choi et al., 2025).

Taken together, on the one hand, astrocytes play a central role in the pathogenesis of neuroinflammation (Ye and Quintana, 2026) by both secreting ECM components and sensing ECM-derived mechanical cues. On the other hand, through the augmented production of ECM constituents, reactive astrocytes contribute to pathological ECM remodeling (Pekny and Pekna, 2016) and glial scar formation, while mechanosensitive ion channels (Chi et al., 2022) enable them to translate these microenvironmental changes into inflammatory biochemical signals.

Nevertheless, the direct role of astrocytic Piezo1 in EAE remains an active area of research. In this manuscript, we explore the tripartite interaction between astrocytic expression of Piezo1, ECM component changes and matrix stiffness in chronic EAE and how this interaction contributes to disease development and severity.

## Materials and methods

### Induction of experimental autoimmune encephalomyelitis (EAE)

*N* = 13 female wild-type (WT) C57BL/6J mice were purchased from Charles River Laboratories (Sulzfeld, Germany) and kept under specific pathogen-free conditions at the Haus für Experimentelle Therapie 4 of the University Hospital Bonn, with unlimited access to standard rodent diet and water. All experiments were approved by the Landesamt für Natur-, Umwelt-, und Verbraucherschutz Nordrhein-Westfalen (LANUV; file 81-02.04.2022.A072) and performed in accordance with the German Animal Welfare Law. They also complied with the “Principles of Laboratory Animal Care” (IACUC guidelines), the 3R principle, and the Animal Research: Reporting of In Vivo Experiments (ARRIVE) guidelines (Kilkenny et al., 2010).

At 8 weeks old, 10 animals were immunized with MOG:_35-55_ (Cat. No. EK-2110, Hooke Laboratories) in complete Freund’s adjuvant (CFA containing 1 mg/mL MOG:_35-55_ and 1-5 mg/mL killed *Mycobacterium tuberculosis* H37Ra). Additionally, 100 ng of pertussis toxin (Ptx) (Cat.No. BT-0105, Hooke Laboratories) was applied on the day of immunization and 24 hours later.

The weight of the mice and clinical signs of disease were monitored daily. Mice were scored using the standard EAE scoring system (Dahl et al., 2025) : 0 – no symptoms; 1 – floppy tail; 2 = partial weakness of the hind limbs; 3 = complete paralysis of the hind limbs; 4 = quadriplegia; 5 = moribund. A group of *N* = 3 naïve mice was used as control. Together the *N* = 16 mice are referred to as cohort 1 throughout the text.

### Perfusion, tissue dissection and paraffin embedding

Mice were killed using CO_2_ and transcardially perfused with PBS before removal of the spinal column for immersion fixation overnight in 4 % para-formaldehyde (PFA) in PBS. The PFA-fixed tissue from the different mice were then processed for paraffin embedding according to a standardized protocol (Lowinski et al., 2025). Paraffin sections were cut at 10 µm thickness and stained for various antigens to assess pathology.

### Immunohistochemistry

Immunohistochemical staining for the different antigens was performed as previously described by our group (Chunder et al., 2022, 2023; Lowinski et al., 2025). Briefly, murine spinal cord sections were deparaffinized with xylene and rehydrated using a descending series of isopropanol. Heat-mediated antigen retrieval was carried out using 10 mM sodium citrate buffer (pH 6.0). All incubation steps were done in a humified chamber and slides were washed with tris-buffered saline (TBS) containing 0.05% Tween-20 (TBS-T) between every incubation.

The slides were blocked with 5% milk powder in TBS-T (blocking buffer) for 1 hour at RT. Primary antibodies were diluted in 0.5% blocking buffer, and sections were incubated with the antibody solution O/N at 4°C. For double stainings, two primary antibodies were mixed into a single cocktail and applied simultaneously onto the slide in a single-reaction step. After washing the slides with TBS-T, the appropriate seconday antibodies, either individually or as a cocktail of two, diluted in 0.5% blocking buffer, were applied to the sections, which were then incubated at RT for 2 hours in the dark. All sections were subsequently stained using Fluoroshield mounting medium containing 42,6-diamidino-2-phenylindole (DAPI) (Abcam, Cambridge, UK). For all the different stainings, a technical negative control was included where the tissue section was incubated with secondary antibodies only.

Details of all the primary and secondary antibodies, including their dilutions, are listed in

**Table 1.**

|  |  |  |  |  |  |
| --- | --- | --- | --- | --- | --- |
| CD3 | rabbit | monoclonal | ab135372 | Abcam | 1:150 |
| Chondroitin sulfate (CS56) | mouse | monoclonal IgM | C8035-.2ML | Merck/Sigma | 1:400 |
| Fibronectin | mouse | monoclonal | 66042-1-Ig | Proteintech | 1:100 |
| GFAP | chicken | poyklonal | ab4674 | Abcam | 1:500 |
| IBA1 | rabbit | polyclonal | 019-19741 | Wako/VWR | 1:500 |
| MMP3 | rabbit | polyclonal | AB53015 | Abcam | 1:100 |
| MMP10 | rabbit | polyclonal | ABT289 | Merck/Millipore | 1:100 |
| OPN | rabbit | polyclonal | AB8448 | Abcam | 1:200 |
| PIEZO1 | rabbit | polyclonal | 15939-1-AP | Proteintech | 1:200 |
| SMI-32 | mouse | monoclonal | ENZ-ABS219-0100 | Enzo | 1:500 |
| SMI-99 | mouse | monoclonal | 808401/100µl | BioLegend | 1:500 |

### Epifluorescence microscopy and confocal imaging

Tile scans covering each spinal cord section were acquired using a DMi8 Thunder imager (Leica) microscope equipped LAS X software (Leica Application Suite X 3.8.2.27713) at different magnifications as indicated in the figure legends.

Confocal images were acquired on an A1-HD25 inverted confocal microscope equipped with a 40x/1.15 NA water immersion objective and an A1 large-field-of-view (LFOV) camera (all from Nikon). Fluorescence channels of each image were captured sequentially. The confocal pinhole size was set to 16.6 µm (1.0 airy unit), resulting in a optical Z slice thickness of 0.37 µm. For GFAP and PIEZO1 colocalization, single confocal slices were captured and post-processed using a customized image analysis macro on Fiji (Nurmatov, Hintze, Kuerten, 2026).

### Cell culture

Mixed glial cultures were generated from cortices of WT C57BL/6J mouse pups between postnatal days 1-3. Cortices from *N* = 4 pups were pooled together for every isolation. After dissection into 1-3 mm pieces in ice-cold dissection medium (DMEM supplemented with penicillin/streptomycin and 1 mM HEPES, pH 7.4; all components were purchased from Gibco), the cortices were digested in 0.25 % trypsin (Cat. No. 15090046, Gibco) for 20 min at 37°C. Trypsination was stopped by 5 washes with pre-warmed dissection medium, followed by tissue dissociation using sterile Pasteur pipettes of two consecutively decreasing tip diameters. After complete dissociation of the tissue, the cell suspension was diluted in 30 mL of DMEM supplemented with 10 % heat-inactivated fetal bovine serum (Cat. No. A5670801, Gibco) (plating medium) and divided into two tissue culture grade T75 flasks (Sarstedt) pre-coated with 0.00075% (w/v) poly-D-lysine (Cat. No. P6407, Merck). Cultures were maintained at 37°C and 5% CO_2_ for two days after which the non-adherent cells were aspirated and the remaining cells were replenished with fresh plating medium. Cells were allowed to reach confluency at 37°C and 5% CO_2_ for a week and were then passaged at a ratio of 1:3. Glial cultures between passages P2 and P5 were used for further experiments.

#### Preparation of hydrogels of different stiffnesses

Hydrogels with tunable stiffness composed of crosslinked oxidized hyaluronic acid (OHA) and gelatin (GEL) were fabricated as previously described (Lorke et al., 2025). All components were dissolved in D-PBS (without Ca^2+^ and Mg^2+^). Briefly, solutions of OHA and microbial transglutaminase (mTG) (SigmaAldrich) were mixed with a GEL solution and transferred into tissue culture grade 6-well or 48-well plates which were pre-coated with 0.00075% (w/v) poly-D-lysine. Hydrogel stiffness was tuned by adjusting the OHA concentration (Table 2). After gelation for 60 min at RT, gels were submerged in DMEM with penicillin/streptomycin for a maximum of 3 days until plating of the mix glial cells.

**Table 2:**
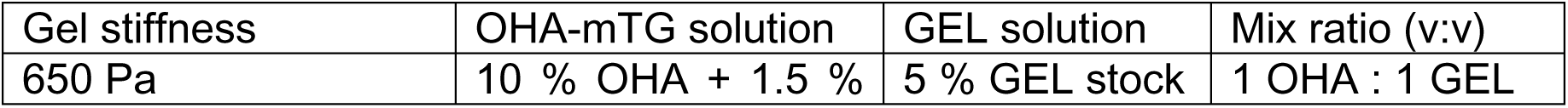
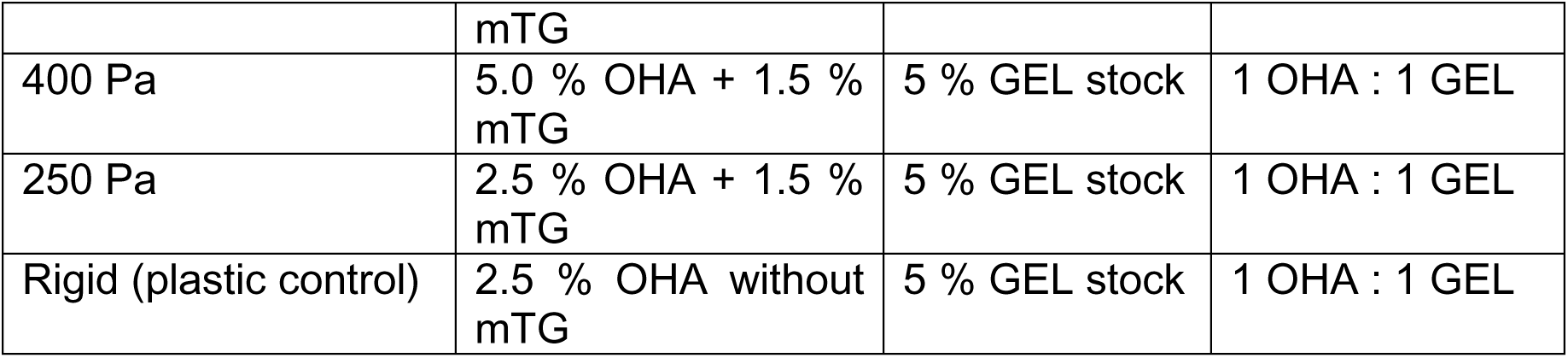
Tunable gel stiffness recipe.

### RNA sequencing of mixed glial cultures

For RNA sequencing, 100,000 cells per well were plated in plating medium either on hydrogel-coated surfaces of varying stiffness or on plastic control surfaces. After 24 hours, plating medium was replaced with assay medium (DMEM supplemented with 0.5 % heat-inactivated fetal bovine serum).

After 48 hours in assay medium, cells were trypsinized with 0.5 % trypsin for 10 minutes at 37°C and collected by centrifugation at 300 g. Cell pellets were snap-frozen in liquid nitrogen and stored at -80°C until RNA extraction. RNA from the cells were collected from two separate preparations of mixed glial cultures plated on hydrogels (stiffnesses: 250 Pa or 650 Pa), or OHA/GEL-coated rigid tissue culture plastic and sent to the Institute of Human Genetics at FAU Erlangen-Nürnberg for bulk RNA sequencing.

### Luciferase assay

For luciferase assays, 35,000 cells in plating medium were disseminated per well of a 48-well plate containing the hydrogels or the plastic control surface and incubated overnight at 37°C and 5% CO_2_. Cells were then transfected using the XFect™ transfection reagent (Takara) according to the manufacturer’s recommendations. Cells were co-transfected with 2.7 µg of the following luciferase reporter/well – YAP/TAZ (Cat. No. 34615, Addgene); NFAT (Cat. No. 17870, Addgene); SRE (Cat. No. pGL4.33, Promega) and 300 ng renilla luciferase (a kind gift from Isabella Graef, Stanford University).

6 hours after transfection, plating medium was replaced with assay medium (DMEM supplemented with 0.5 % heat-inactivated fetal bovine serum) with or without additional drugs (5 µM Yoda1 or 5 µM GsMTx4) as indicated in Figure 5A. 48 hours later, the supernatants were collected and cells were washed in PBS before lysing them in 100 µL passive lysis buffer (Promega) for 20 min at RT. 50 µL of the cell lysate were transferred to an opaque white 96-well plate to measure firefly and renilla luciferase activities in a plate reader (Corning). For each sample, firefly activity was normalized to renilla activity and all values of a given reporter construct were normalized within each experiment to the corresponding rigid plastic surface control. All luciferase experiments were performed at least 3 times. Each data point corresponds to one individually transfected sample well.

### Statistical analysis

Data were statistically analyzed in Prism Version 9.5.1 (Graph Pad). For imaging and clinical data, non-normal distribution was assumed for 2-way ANOVA with a two-stage linear step-up procedure of Benjamini, Krieger and Yekutieli. Spearman correlation coefficients r were calculated and tested with a two-tailed exact P value approximation. For luciferase reporter assays, 2-way ANOVA with Tukey’s multiple comparison test was performed.

## Results

### Extracellular matrix remodeling shows regional differences in the spinal cord during EAE

To investigate how remodeling of the extracellular matrix is linked to CNS tissue stiffness changes associated with EAE *in vivo*, we performed immunofluorescent staining on spinal cord (SC) cross sections of mice with acute and chronic EAE. We induced EAE by immunization of 10 female C57BL/6J wildtype mice (8 weeks old) with the myelin oligodendrocyte glycoprotein peptide (MOG_35-55_), and sacrificed 5 mice 17 days post immunization (17 dpi, acute EAE; mean EAE score ± SEM 2.3 ± 0.27) and 5 mice at 31 dpi (chronic EAE; mean EAE score ± SEM 1.55 ± 0.55) (Fig. 1A).

**Figure 1.**
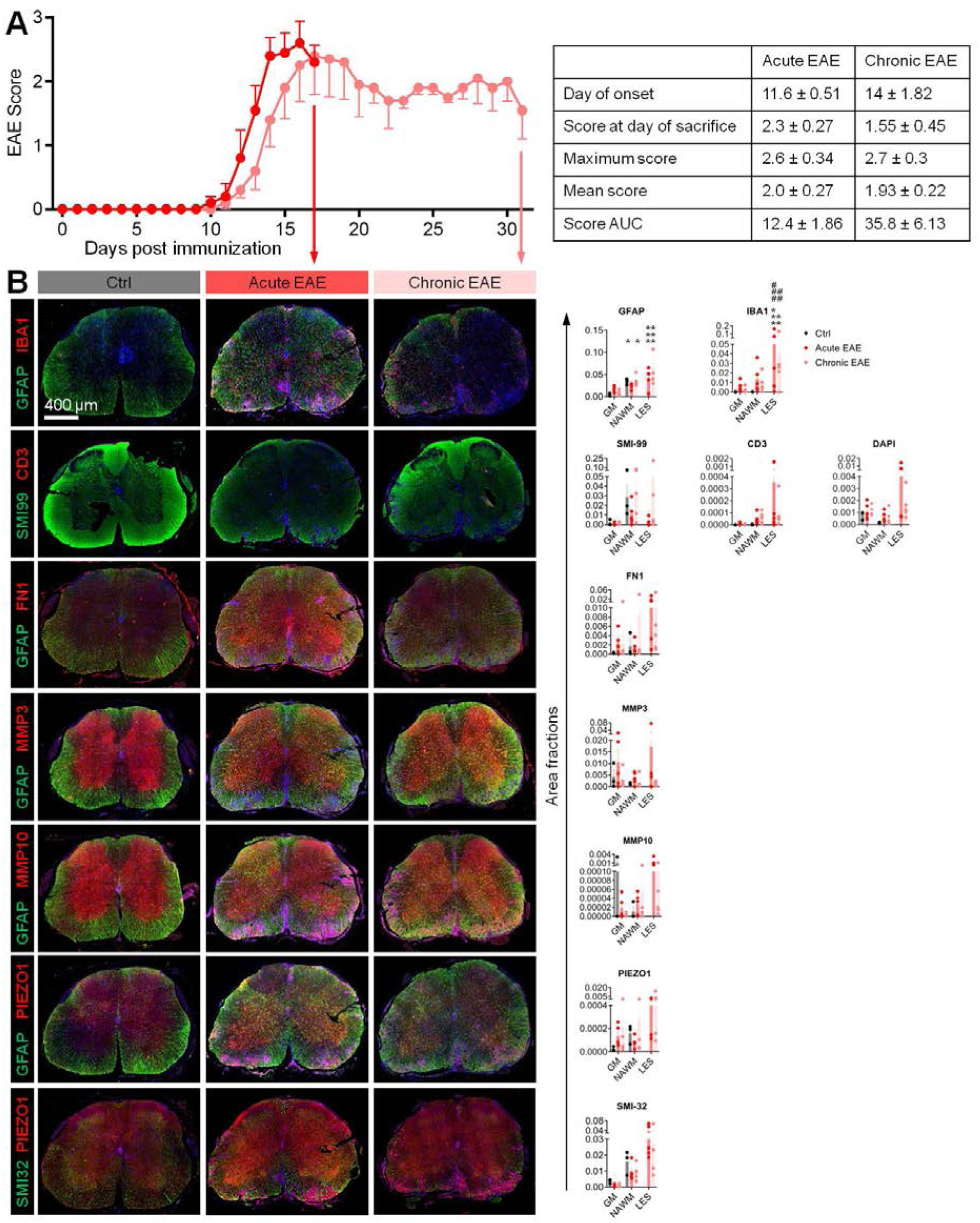
Components of extracellular matrix remodeling are regionally upregulated in the spinal cord during EAE. **(A)** Clinical disease course of MOG_35–55_-induced EAE in C57BL/6 mice. Animals were sacrificed during acute EAE (17 post-immunization) or chronic EAE (day 31 post-immunization). Clinical parameters for both cohorts are summarized in the table. **(B)** Representative spinal cord cross-sections from control mice and mice with acute or chronic EAE stained for markers of astrocyte activation (GFAP), immune cell infiltration (IBA1, CD3), axonal integrity (SMI-99, SMI-32), extracellular matrix remodeling (FN1, MMP3, MMP10), and mechanosensory signaling (PIEZO1). Nuclei were counterstained with DAPI. Quantification of marker-positive area fractions in gray matter (GM), normal-appearing white matter (NAWM), and lesion (LES) regions is shown on the right. Individual data points represent single animals; bars indicate mean ± SEM. Increased expression of GFAP, FN1, MMP3, MMP10, and PIEZO1 was observed in a region-dependent manner during EAE, with the most pronounced changes detected in lesion areas. Statistical comparison vs. respective GM Ctrl: * p < 0.05, ** p < 0.01, *** p < 0.005; Statistical comparison vs. respective NAWM Ctrl: ## p < 0.01, ### p < 0.005

We next performed region-specific quantification of EAE-induced histopathology markers which indicated distinct regulation between gray matter, normal-appearing white matter and lesion areas (GM, NAWM, and LES, respectively) during both acute and chronic EAE phases (Fig. 1B). The myelin marker SMI-99 (against myelin basic protein, MBP) was strongly decreased in LES and mildly decreased in acute and chronic EAE NAWM. Immune cell markers IBA1 and CD3 were highest during acute and chronic EAE in the LES areas. In the NAWM, both markers were increased in acute EAE but reverted somewhat during chronic EAE. In the GM, IBA1 was strongly upregulated during acute EAE and remained slighty up during chronic EAE, while CD3 remained virtually unchanged. The high levels of DAPI signal in the acute EAE LES indicate high cell densities which were less noticeable in the GM and NAWM during EAE. Thus, T cell (and likely, other immune cell) infiltration stays grossly confined to the WM lesions without spillover into the NAWM or GM. Nevertheless, IBA1 upregulation in the GM indicates microglia activation in the GM despite the absence of noticeable immune cell infiltration into this region.

ECM in the CNS is mainly synthesized and regulated by astrocytes, the major non-neuronal parenchymal cell type in the brain and spinal cord. We stained lumbar spinal cord cross sections for the astrocyte marker GFAP (glial fibrillary acidic protein) in combination with the ECM molecules fibronectin (FN1), matrix metalloproteinases 3 and 10 (MMP3 and MMP10, respectively) to reflect ECM structural and remodeling components. In healthy controls (Ctrl), the GFAP signal in NAWM was higher than in GM. During acute EAE, GFAP was strongly upregulated in GM and remained increased to a lesser extent during chronic EAE. In WM regions, GFAP was only slightly increased in the LES but remained unchanged in NAWM compared to Ctrl WM. The ECM molecules FN1, MMP3 and MMP10 were all detected at highest levels in the acute EAE LES regions, but FN1 and MMP3 were also upregulated in NAWM and GM during acute EAE.

Because ECM remodeling might induce tissue stiffness changes during EAE and potentially triggers cellular mechanosensory signaling pathways, we also stained for the *bona fide* mechanosensory ion channel PIEZO1 which is expressed on various cell types in the CNS, including neurons and all glial cell types (Silva et al., 2024). PIEZO1 expression was highest inside the LES regions during acute and chronic EAE, reflecting the high expression of PIEZO1 on leukocytes. In healthy controls, the PIEZO1 signal in NAWM was higher than in GM. While GM PIEZO1 levels were upregulated during acute EAE and returned to almost normal levels in chronic EAE, PIEZO1 in NAWM remained unchanged throughout the course of EAE. Thus, changes of PIEZO1 levels parallelled the pattern of changes observed for GFAP in different SC regions and throughout the course of EAE.

Taken together, the astrocyte marker GFAP, the ECM constituents FN1, MMP3 and MMP10, and the mechanosensory ion channel PIEZO1 all showed parallel patterns of regional changes during EAE with highest levels in LES throughout the disease, and temporary upregulation in the GM during acute EAE, while NAWM remained relatively unaffected.

#### The mechanosensory ion channel PIEZO1 is upregulated on spinal cord astrocytes during EAE

To further investigate the parallel regional regulation of PIEZO1 and GFAP expression in different SC regions throughout EAE, we performed overlap analysis on confocal images (Fig. 2A). Again, the transient upregulation of total PIEZO1 in GM in acute EAE was slightly attenuated during chronic EAE, but remained elevated compared to control levels (Fig. 2B, red bars in left panel). When we analyzed astrocyte-specific PIEZO1, we observed that its expression remained consistently elevated in GM during both acute and chronic EAE (Fig. 2B, yellow bars in the right panel). In NAWM and LES areas, both total PIEZO1 and astrocytic PIEZO1 levels were upregulated during acute EAE and increased further during chronic EAE.

**Figure 2.**
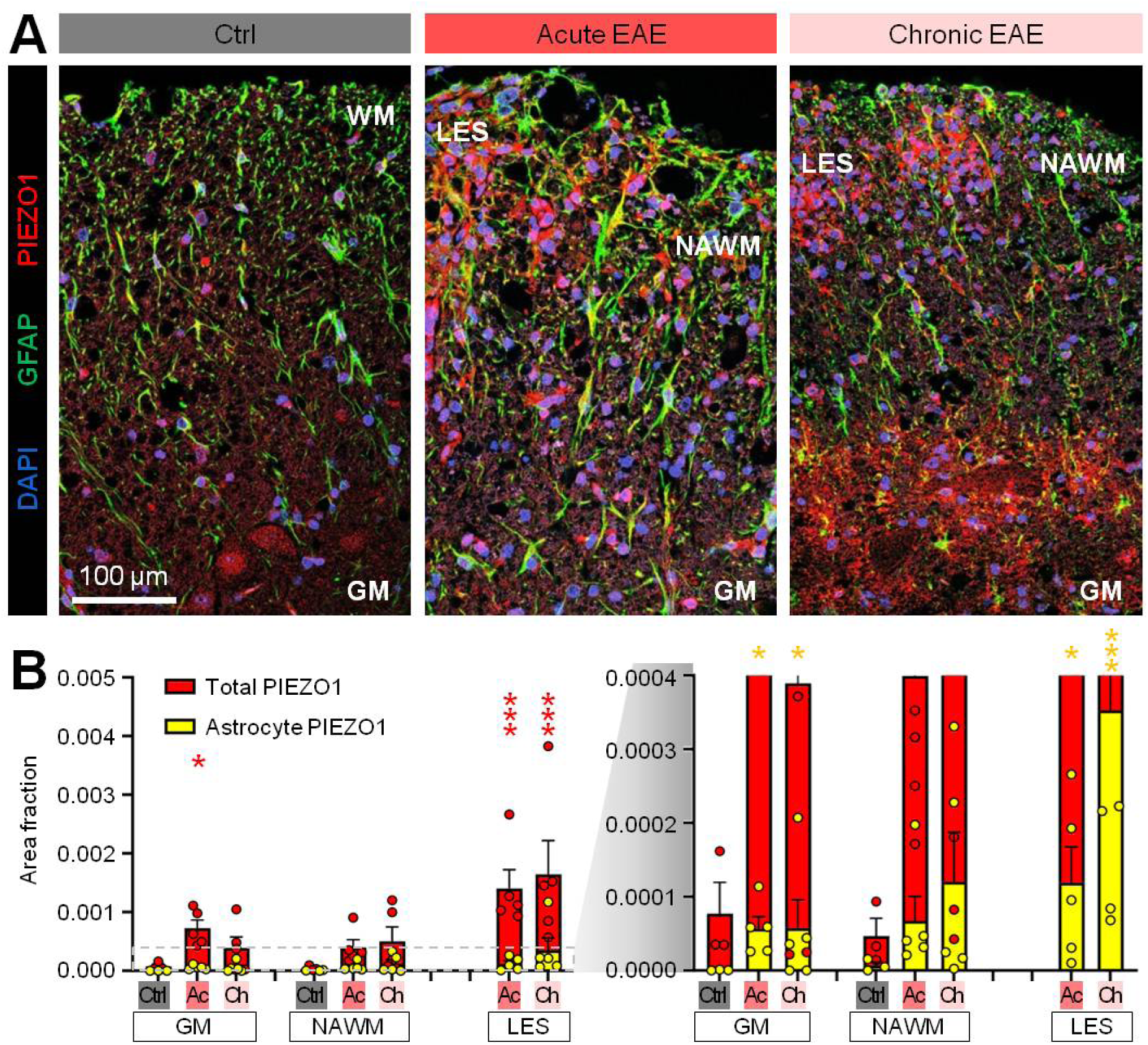
The mechanosensory ion channel PIEZO1 is upregulated in spinal cord astrocytes during EAE. **(A)** Representative immunofluorescence images of spinal cord sections from control mice and mice with acute or chronic EAE stained for GFAP (green), PIEZO1 (red), and DAPI (blue). Gray matter (GM), normal-appearing white matter (NAWM), and lesion (LES) regions are indicated as identified on characteristic GFAP and DAPI staining patterns. **(B)** Quantification of total PIEZO1-positive area fractions (red) and astrocyte-associated PIEZO1-positive area fractions (yellow; defined as PIEZO1 signal colocalizing with GFAP) in GM, NAWM, and LES regions. Individual data points represent single animals, and bars indicate mean ± SEM. Astrocytic PIEZO1 expression was elevated during both acute and chronic EAE, whereas total PIEZO1 expression displayed region-specific changes, with the most pronounced increases observed in lesion areas. Dashed lines indicate magnified area of the chart as represented in the right chart. GM, gray matter; NAWM, normal-appearing white matter; LES, lesion area. Statistical comparison of PIEZO1 levels vs. respective GM Ctrl: * p < 0.05, *** p < 0.005. Red asterisks indicate statistical comparison of total PIEZO1 (left bar chart), yellow asterisks indicate statistical comparison of astrocyte-specific PIEZO1 (right bar chart).

These findings suggest that, although astrocytic PIEZO1 represents only a minor fraction of total PIEZO1 expression in the SC, it contributes substantially to the overall increase in PIEZO1 associated with non-immune cells.

#### Clinical EAE correlates with PIEZO1 expression in GM and NAWM

Based on the concomitant upregulation of the astrocyte marker GFAP and PIEZO1 and the known association of GFAP upregulation with astrogliosis, we tested correlations between local PIEZO1 expression changes and clinical EAE parameters. We correlated the EAE score at the time of sacrifice (endpoint EAE score), total EAE load (EAE – AUC, area under the curve), maximum EAE score as well as the weight at the time of sacrifice (endpoint weight) with the histological quantification of immune pathology markers, ECM markers, and PIEZO1 levels (Fig. 3A). We identified a significant regional correlation of the

**Figure 3.**
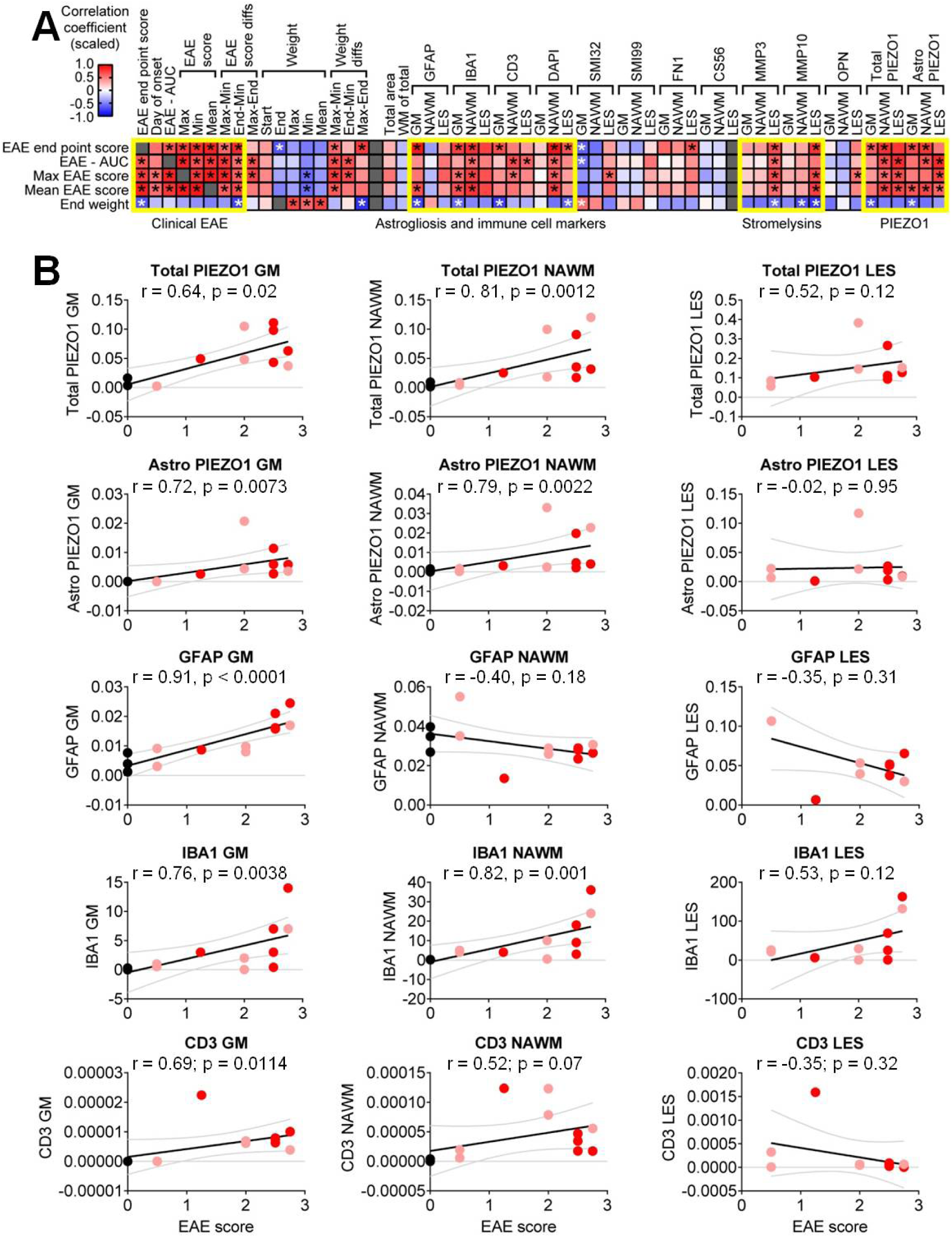
Astrocytic PIEZO1 upregulation is regionally correlated with clinical EAE severity. **(A)** Correlation matrix summarizing Spearman correlation coefficients between clinical EAE parameters (EAE score at sacrifice, maximum score, mean score, score area under the curve [AUC], and weight), histopathological markers, extracellular matrix remodeling markers, and PIEZO1 expression in gray matter (GM), normal-appearing white matter (NAWM), and lesion (LES) regions. Color intensities are scaled to match maximum and minimum values. **(B)** Representative correlations between clinical EAE score and total PIEZO1, astrocyte-associated PIEZO1, GFAP, IBA1, and CD3 expression in GM, NAWM, and LES regions. Each data point represents one animal. Lines indicate linear regression fits with 95% confidence intervals. Spearman correlation coefficients (r) and corresponding p-values are indicated in each panel. Among the analyzed markers, GFAP expression in GM showed the strongest correlation with EAE score, while GFAP in NAWM or LES regions was not correlated with the EAE score. Both total and astrocytic PIEZO1 expression in GM and NAWM, but not in LES, correlated positively with clinical EAE score.

EAE endpoint score with GFAP in GM, but not in NAWM or LES. The three ECM components FN1, MMP3 and MMP10 were significantly correlated with the EAE endpoint score only in the LES, but not in other regions. Total and astrocytic PIEZO1 levels were both correlated with the EAE endpoint score in GM and NAWM, but not in the LES.

Among all markers analyzed, GFAP levels in GM showed the strongest correlation with clinical EAE severity (Fig. 3B). Notably, both total and astrocyte-specific PIEZO1 levels in GM and NAWM also exhibited strong correlations with clinical disease parameters, surpassing those observed for the classical inflammatory markers IBA1 and CD3, which are commonly regarded as robust indicators of disease activity. In LES regions, none of the analyzed markers correlated with clinical EAE parameters.

#### Locally altered PIEZO1 levels in EAE correlate with pathology and ECM markers in distinct SC areas

The most prominent histopathological changes during acute and chronic EAE are commonly believed to occur mainly within the LES areas. We thus asked whether altered PIEZO1 levels might be locally correlated with histopathology or ECM markers (Fig. 4A). The increased PIEZO1 levels in GM and NAWM were mutually correlated with GFAP, IBA1 and CD3, but the same correlations were not observed inside the LES. By contrast, local increases of PIEZO1 levels in GM and NAWM were correlated with MMP3 and MMP10 levels in the LES, indicative of non-local correlations between these parameters. A more detailed correlation analysis of the GM astrocyte PIEZO1 levels with total PIEZO1, GFAP and ECM markers (Fig. 4B) showed that altered astrocyte PIEZO1 levels within the GM were correlated with MMP10 expression inside the LES, but not locally within the GM.

**Figure 4.**
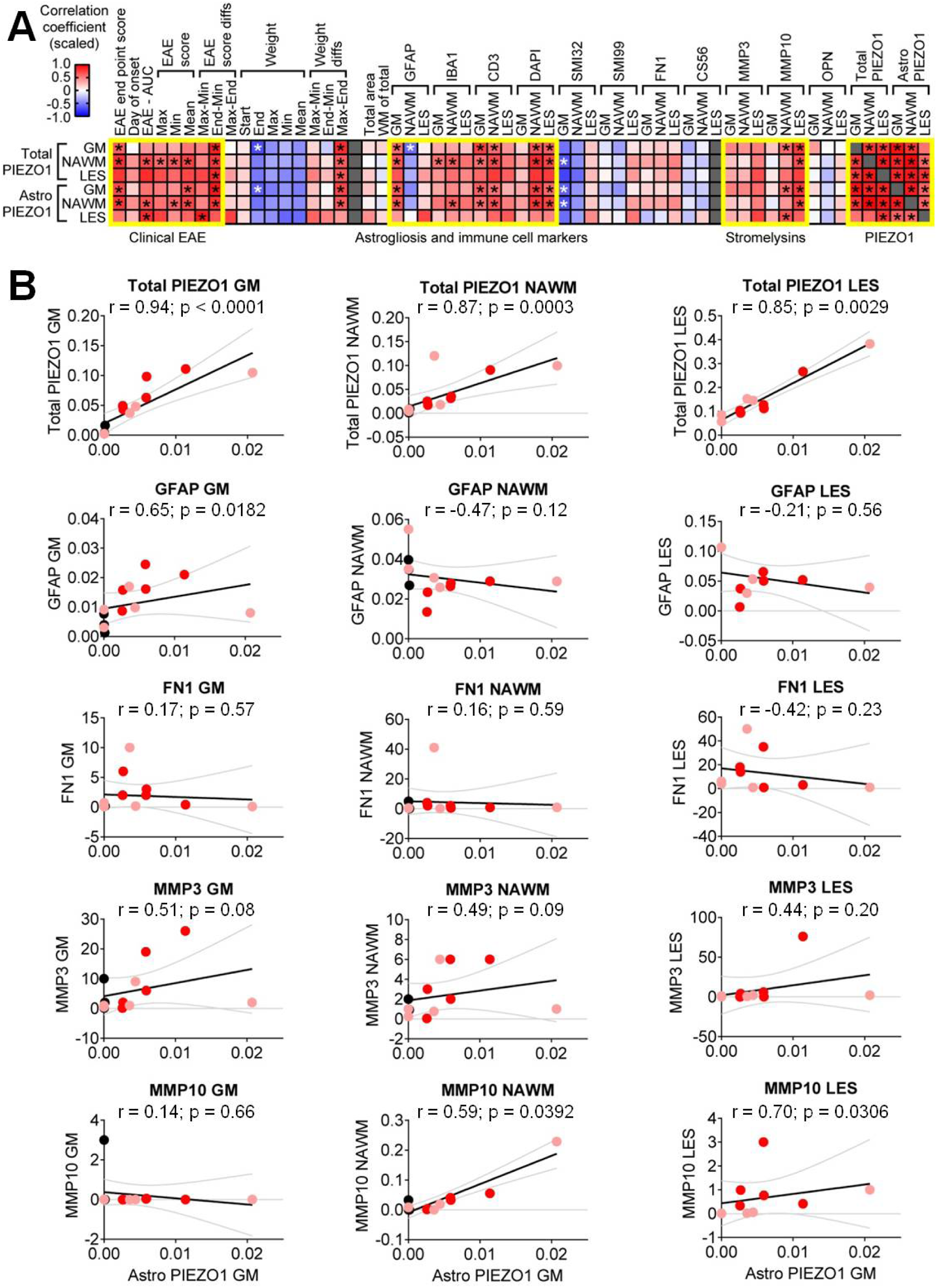
Astrocytic PIEZO1 upregulation is regionally associated with extracellular matrix remodeling during EAE. **(A)** Correlation matrix summarizing Spearman correlation coefficients between GM astro-cyte-associated PIEZO1 expression and clinical EAE parameters, markers of astrogliosis and immune cell infiltration, and extracellular matrix remodeling markers in gray matter (GM), normal-appearing white matter (NAWM), and lesion (LES) regions. Color intensities are scaled to match maximum and minimum values. **(B)** Representative correlations between astrocyte-associated PIEZO1 expression and to-tal PIEZO1, GFAP, FN1, MMP3, and MMP10 expression across GM, NAWM, and LES re-gions. Each data point represents one animal. Lines indicate linear regression fits with 95% confidence intervals. Spearman correlation coefficients (r) and corresponding p-values are shown in each panel. GM astrocytic PIEZO1 expression showed strong correlations with total PIEZO1 expres-sion across all analyzed regions. Correlations of GM astrocytic PIEZO1 with markers of extracellular matrix remodeling were not confined to intra-regional correlations but exhibit-ed cross-correlation between GM and LES regions. GM, gray matter; NAWM, normal-appearing white matter; LES, lesion area.

**Figure 5.**
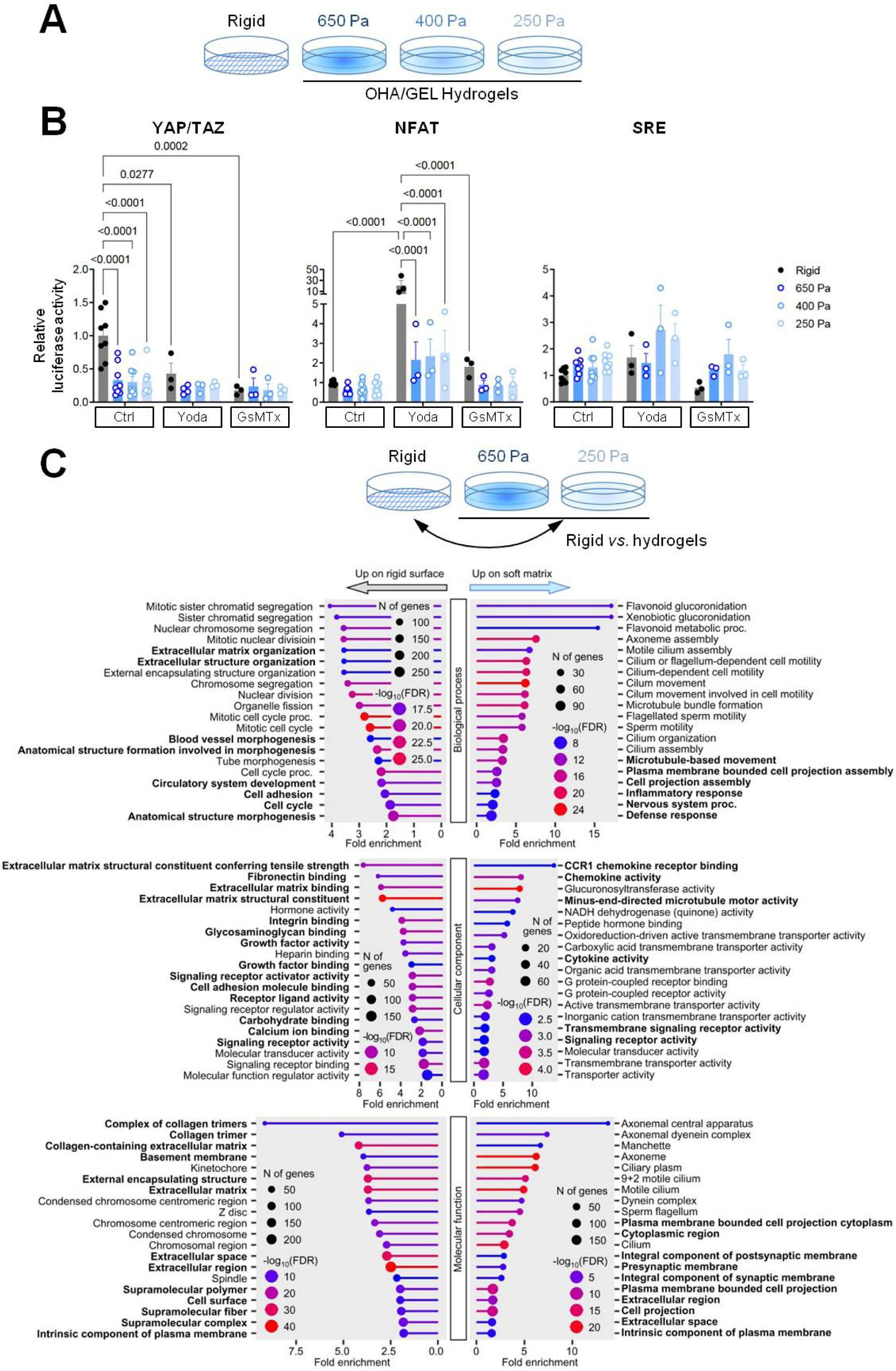
Environmental stiffness induces extracellular matrix- and cytokine-associated transcriptional programs in primary glial cells. **(A)** Experimental design. Mixed primary murine glial cells were cultured either on rigid pol-ystyrene surfaces or on oxidized hyaluronic acid-gelatin (OHA/GEL) hydrogels with de-fined elastic moduli (650 Pa, 400 Pa, and 250 Pa) to model different mechanical microen-vironments. **(B)** Activity of mechanosensitive luciferase reporters in glial cells cultured on substrates of varying stiffness. Relative activities of YAP/TAZ-, NFAT-, and serum response element (SRE)-associated signaling pathways were quantified under the indicated culture condi-tions. Data are presented as individual samples and mean ± SEM. Statistical significance (non-parametric ANOVA) is indicated in the respective panels. **(C)** Gene ontology (GO) enrichment analysis comparing glial cells cultured on rigid sub-strate and compliant hydrogels. Differentially expressed genes were analyzed for enrich-ment of biological process, molecular function, and cellular component GO terms. Cells cultured on rigid substrates preferentially upregulated pathways associated with cell cycle progression, extracellular matrix organization, cell adhesion, and growth factor signaling, whereas cells cultured on softer matrices showed enrichment of cytokine-related, inflam-matory, and cytoskeletal pathways. Dot size indicates the number of genes associated with each GO term, and color intensity reflects statistical significance.

#### Environmental stiffness induces ECM and cytokine gene expression in astrocytes *in vitro*

To test whether stiffness changes of the extracellular environment can induce pathologically relevant changes of cellular gene expression, we plated neonatal mouse astrocytes on rigid plastic surface and hydrogels composed of enzymatically cross-linked oxidized hyaluronic acid and gelatin (OHA/GEL) with tunable stiffness between 250 Pa and 650 Pa (Fig. 5A) (Lorke et al., 2025). We first confirmed mechanosensation in these cells by luciferase reporters (Fig. 5B). The mechanosensory YAP/TAZ pathway luciferase reporter was reduced to less than 50 % compared to the rigid plastic surface. Application of the PIEZO1 agonist Yoda1 significantly reduced YAP/TAZ reporter activity on the rigid plastic surface but had no effect on the hydrogels. Likewise, the PIEZO1 antagonist GsMTx4 significantly reduced YAP/TAZ reporter activity on plastic but remained ineffective in the hydrogels. By contrast, the Ca^2+^-dependent NFAT pathway luciferase reporter remained unchanged between plastic and hydrogel conditions in the absence of Yoda1 or GsMTx4. Addition of Yoda1 significantly increased NFAT reporter levels on rigid plastic to over 20-fold of untreated rigid plastic conditions, and this Yoda1-induced high reporter activity was almost 10-fold reduced on the soft hydrogels. Treatment with the PIEZO1 inhibitor GsMTx4 had no significant effect on NFAT luciferase reporter levels. As a control, we tested the unrelated serum response element (SRE) luciferase reporter which showed no major differences between stiffness conditions or pharmacological activation or inhibition of PIEZO1 activity.

Having showed mechanosensitive responsiveness of mouse astrocytes on hydrogels versus rigid plastic, we subjected cells grown on rigid plastic or hydrogels to bulk RNA-seq analysis. Differentially expressed genes (DEGs) were identified as genes that were upregulated by 2-fold or downregulated by 0.5-fold on hydrogels in comparison to rigid plastic surface. DEG lists were then analyzed by gene ontology (GO) term analysis (Fig. 5C) using ShinyGO 0.80 (Ge et al., 2020). GO term analysis showed that on rigid plastic surface, astrocytes upregulated ECM-related terms, cell adhesion terms as well as cell signaling and blood vessel development-related terms. By contrast, on soft hydrogels upregulated terms frequently related to cytoskeletal organization, cell motility and cell projection, but also inflammation and cytokine signaling.

## Discussion

The central aim of this study was to investigate whether neuroinflammatory tissue remodeling during EAE is associated with activation of mechanosensory pathways in the CNS, and how long-ranging the ensuing effects are.

In a first set of experiments, our analyses revealed pronounced regional differences in ECM remodeling during EAE. While inflammatory infiltrates remained largely confined to demyelinated white matter lesions, increased expression of GFAP, FN1, and MMP3 was also detected in gray matter regions, indicating that glial-driven tissue remodeling extends beyond areas of overt immune cell accumulation. These findings are consistent with the established role of reactive astrocytes as key regulators of ECM composition during neuroinflammation (Lau et al., 2013; Harlow and Macklin, 2014).

A notable observation was the close correspondence between the regional expression patterns of GFAP and the mechanosensitive ion channel PIEZO1. Similar to astrocyte activation, total PIEZO1 expression was strongly increased within lesions and transiently elevated in gray matter during acute EAE, whereas NAWM remained comparatively unaffected. Given that ECM composition is a major determinant of tissue mechanical properties, these findings suggest that inflammatory ECM remodeling and mechanosensory signaling may be functionally linked during EAE. Along these lines, recent studies have identified PIEZO1 as a regulator of glial and immune cell responses during neuroinflammation (Velasco-Estevez et al., 2018; Atcha et al., 2021).

Although the present data do not establish a causal relationship between ECM remodeling and PIEZO1 regulation, the parallel changes observed across spinal cord regions support the concept that mechanosensory pathways are engaged during neuroinflammatory tissue remodeling.

To further explore the relationship between mechanosensory signaling and neuroinflammatory tissue remodeling, we next focused on the cellular expression pattern of PIEZO1 during EAE. Upregulation of astrocytic PIEZO1 during both acute and chronic EAE supports the concept that astrocytes actively sense and respond to mechanical changes in the neuroinflammatory microenvironment (Keating and Cullen, 2021). Given that reactive astrocytes contribute substantially to extracellular matrix remodeling (Woo and Sontheimer, 2023), increased Piezo1 expression may establish a self-feeding mechanotransduction loop that promotes persistent astrogliosis (Wan et al., 2023). The elevation of astrocytic PIEZO1 in NAWM further suggests that mechanical signaling may accompany early tissue remodeling events even before lesion development. This is of particular importance in MS as NAWM has been long recognized as not being truly normal but rather undergoing subtle inflammatory changes (Allen, McQuaid, Mirakhur and Nevin, 2001) and possibly experiencing diffuse tissue remodeling (de Groot et al., 2013).

Furthermore, our findings also support the notion that astrocytic PIEZO1 acts as a mechanosensitive consolidator of neuroinflammatory tissue remodeling. PIEZO1 expression in the GM and NAWM appears to capture diffuse pathological changes (Gonzales et al., 2025) that positively correlate with disease severity.

The observation that PIEZO1 expression in GM and NAWM correlated with lesion-associated MMP3 and MMP10 levels supports a role for extracellular matrix remodeling (Jetta et al., 2021) in driving mechanosensitive responses during EAE. Matrix metalloproteinases are major mediators of ECM degradation and restructuring (Wan et al., 2021) and contribute to alterations in tissue stiffness and architecture (Xie et al., 2014) during neuroinflammation. As PIEZO1 functions as a mechanosensitive ion channel that responds to physical changes in the cellular microenvironment, increased PIEZO1 expression may represent an adaptive response to ECM remodeling occurring during disease progression.

The strong association between astrocytic PIEZO1 expression and disease severity raises the question of whether changes in the physical tissue environment can directly influence glial cell behavior. To address the potential biological consequences of altered tissue mechanics, we investigated the transcriptional responses of primary murine astrocytes exposed to substrates of different stiffness *in vitro*.

Gene ontology analysis of mixed primary murine glial cells cultured either on polystyrene surface or on oxidized hyaluronic acid revealed a profound stiffness-dependent transcriptional response. Cells cultured on rigid surface (as shown in Fig. 5C), with an estimated Young’s modulus that is several thousand fold higher (Acevedo-Acevedo and Crone, 2015) than CNS tissue under physiological conditions (Weickenmeier et al., 2016), upregulated pathways associated with mitosis, extracellular matrix organization, cell adhesion, and growth-factor signaling. These findings suggest that an increased matrix stiffness drives glial cells toward a reactive phenotype (Williamson et al., 2021) characterized by enhanced proliferation and extracellular matrix remodeling. In contrast, glial cells cultured on softer matrices preferentially expressed genes involved with cytokine activity, inflammatory response and cytoskeleton reorganization which are also hallmarks of a reactive cellular phenotype (Ren et al., 2010). While one study has reported that astrocytes adopt a reactive and proliferative phenotype on sensing a softer environment (Benincasa et al., 2024), another group (Wilson, Hayward and Kidambi, 2016) has shown that astrocytes grown on stiff substrates displayed an astrogliosis-like morphology. Therefore, whether astrocytes switch to a pro-inflammatory or regulatory phenotype on sensing a stiff or soft enviroment is highly context dependent, especially given that 2D culturing of glial cells do not accurately capture their physiological responses and behaviors (Greaney et al., 2025). Nevertheless, our data indicate that astrocytes strongly respond to matrix stiffness changes and astrocyte reactivity can be mechanobiologically modulated to control inflammation or accelarate tissue healing (Cieri and Ramos, 2024; Katiyar et al., 2016).

Taken together, our findings indicate that astrocyte-mediated mechanical signaling may prominently contribute to neuroinflammation and neurological dysfunction in EAE. We identify upregulation of astrocytic PIEZO1 as a prominent component of the tissue response to EAE that is closely associated with extracellular matrix remodeling and clinical disease severity. While the present study does not establish a causal role for PIEZO1 in disease pathogenesis, it highlights mechanosensation as a previously underappreciated aspect of neuroinflammatory tissue remodelling, even in areas outside of the lesion regions directly affected by immune infiltration. Future studies should determine whether PIEZO1 actively shapes astrocyte function and inflammatory responses *in vivo* and assess its potential as a biomarker of disease activity especially in the human disease.

## Authors’ contributions

AB, research, manuscript: review

MH, conceptualization, research, data analysis, manuscript: initial draft, manuscript: review ML, research

MS, ZN, JB, KLH, EB, AE, data analysis

RC, research, data analysis, manuscript: initial draft, manuscript: review SK, conceptualization, research, manuscript: initial draft, manuscript: review

## Acknowledgements

We would like to thank Ji-Young Chang for administrative assistance and Birgit Blanck as well as Marion Michels for technical assistance.

